# PPRD: a comprehensive online database for expression analysis of ~45,000 plant public RNA-Seq libraries

**DOI:** 10.1101/2022.01.28.477949

**Authors:** Yiming Yu, Hong Zhang, Yanping Long, Yi Shu, Jixian Zhai

## Abstract

High-throughput RNA-sequencing (RNA-seq) has become the most popular technology for profiling gene expression in the last decade due to its low cost and high coverage. As a result, the number of RNA-seq libraries from the plant community has been increasing exponentially in recent years. For major crops, such as maize, rice, soybean, wheat, and cotton, the plant community has collected a total of ~45,000 libraries by 2021. To take full advantage of the bigdata of RNA-seq libraries, an effort to integrate all publicly available libraries via a uniformed processing pipeline and curate them into an easy-to-use searchable database is urgently needed. To address this challenge, here we present a comprehensive web-based platform, Plant Public RNA-seq Database (PPRD, http://ipf.sustech.edu.cn/pub/plantrna/). PPRD consists of a large number of RNA-seq libraries of maize (11,726), rice (19,664), soybean (4,085), wheat (5,816), and cotton (3,483) from Gene Expression Omnibus (GEO), Sequence Read Archive (SRA), European Nucleotide Archive (ENA), and DNA Data Bank of Japan (DDBJ) databases.

---

High-throughput RNA-sequencing (RNA-seq) has become the most popular technology for profiling gene expression in the last decade due to its low cost and high coverage. As a result, the number of RNA-seq libraries from the plant community has been increasing exponentially in recent years (Figure 1A). For major crops, such as maize, rice, soybean, wheat, and cotton, the plant community has collected a total of ~45,000 libraries by 2021 (Figure 1B). Although currently there are several RNA-seq databases for plants, for example, CoNekT with 750 rice and 574 maize RNA-seq libraries (Proost and Mutwil, 2018). However, these existing databases only host the already processed data from each study separately, and therefore the expression values cannot be directly compared among projects, because they were derived from different bioinformatic pipelines and often mapped to different versions of the reference genomes. To take full advantage of the bigdata of RNA-seq libraries, an effort to integrate all publicly available libraries via a uniformed processing pipeline and curate them into an easy-to-use searchable database is urgently needed. To address this challenge, here we present a comprehensive web-based platform, <u>P</u>lant <u>P</u>ublic <u>R</u>NA-seq <u>D</u>atabase (PPRD, http://ipf.sustech.edu.cn/pub/plantrna/). PPRD consists of a large number of RNA-seq libraries of maize (11,726), rice (19,664), soybean (4,085), wheat (5,816), and cotton (3,483) from Gene Expression Omnibus (GEO), Sequence Read Archive (SRA), European Nucleotide Archive (ENA), and DNA Data Bank of Japan (DDBJ) databases (Figure 1B). These RNA-seq data are manually curated to highlight different mutants, tissues, developmental stages, abiotic or biotic stresses. Besides showing expression patterns from different tissues and developmental stages (Figure 1C–1E), we also annotated the mutant-related groups and treatment-associated groups in our maize, rice, soybean, wheat, and cotton database, respectively (Figure 1F and 1G). To reduce the quantification biases derived from differing bioinformatic processes, we processed the data of each species with a unified pipeline and the most up-to-date reference genomes (More details on the “Tutorials” page and Supplementary material). Moreover, the database also provided hyperlinks to check the expression level of the homologous genes in other plants and supported a built-in online Integrative Genomics Viewer (IGV) (Figure 1H) (Robinson et al., 2017).

**Figure 1.**
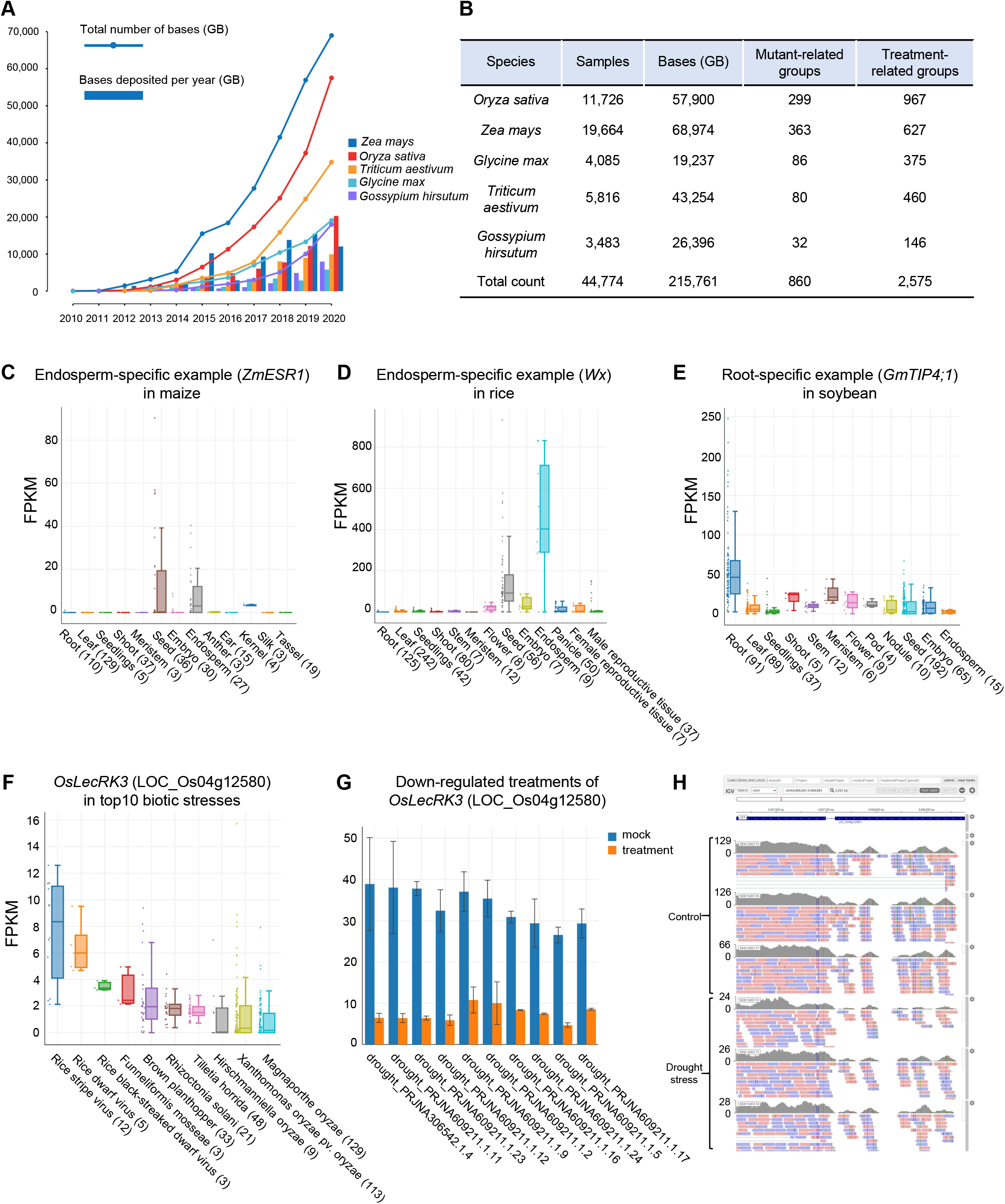
Overview of Plant Public RNA-Seq Database. **(A)** The number of *Oryza sativa, Zea mays, Glycine max, Triticum aestivum*, and *Gossypium hirsutum* sequenced bases per year from 2010 to 2020. Bar indicates the bases deposited per year (GB). Line indicates the total number of bases (GB). GB: giga base pairs. **(B)** The basic summary of RNA-seq libraries. “Mutant-related groups” and “Treatment-related groups” denote the number of groups used to analyze the differential expression. **(C-E)** The tissue-specific expression of some marker genes. The left panel shows the endosperm-specific expression of *ZmESR1* in maize (C), the middle panel shows the endosperm-specific expression of *Wx* in rice (D), and the right panel displays the root-specific expression of *GmTIP4;1* in soybean (E). **(F)** The expression level of *OsLecRK3* (LOC_Os04g12580) among top10 biotic stresses in rice. **(G)** Down-regulated expression of *OsLecRK3* (LOC_Os04g12580) among top10 treatment groups in rice. **(H)** The overview of IGV. The mapped reads of *OsLecRK3* show decreased abundance in drought stress-related samples.

In general, PPRD supports searches by gene ID, library ID, BioProject IDs, keywords, or any combination of these terms in selected libraries. After querying the above terms, the results in tables and diagrams will be returned. Here, we take the query results of a key regulator of plant small RNA biogenesis, *OsDCL3a* (LOC_Os01g68120) (Wei et al., 2014), to illustrate the database. After entering “LOC_Os01g68120” in a “Google-like” search box, the “Information” page will return the basic information of this gene. PPRD also provides hyperlinks for easy access to more information about the corresponding gene in the species-related websites, such as MaizeGDB for maize (Portwood et al., 2019), RGAP for rice (Kawahara et al., 2013), and SoyBase for soybean (Brown et al., 2021). On the “Data Table” page, detailed information could be displayed in a table, and various “Filter” options are designed to allow users to select specific libraries. The “Data Plot” page shows the results of expression comparison in multiple interactive diagrams, including expression levels among different tissues, developmental stages, abiotic and biotic stresses, and up-regulated or down-regulated expression in mutant-related or treatment-related samples. The “CoExpression” page provides a list of genes co-expressed with the searched one. And the “IGV Online” page is flexible for visualizing the mapping landscape of the local genomic region in selected libraries. In addition, the “Share” function was supported to facilitate showing the results with others. Here, we used the tissue-specific expressed genes to validate the results. The expression levels of these genes are consistent with previous studies, such as endosperm-specific expression of gene *ZmESR1* (Zm00001d027820) in maize (Opsahl-Ferstad et al., 1997), endosperm-specific expression of gene *Wx* (LOC_Os06g04200) in rice (Sano, 1984), and root-specific expression of gene *GmTIP4;1* (Glyma.06G084600) in soybean (Song et al., 2016) (Figure 1C–1E).

PPRD also supports users to perform data mining from the large-scale database efficiently. The brown planthopper (BPH) is the most destructive pest that has a massive impact on rice production by the transformations of viruses, and *OsLecRK3* (LOC_Os04g12580) is a crucial gene that confers resistance to the BPH (Liu et al., 2015). As expected, *OsLecRK3* showed higher expression in some viruses-related libraries (Figure 1F). To our surprise, *OsLecRK3* is down-regulated in many drought-related libraries, suggesting that *OsLecRK3* plays a crucial role in drought resistance (Figure 1G). In addition, the mapping details of this gene can be visualized using the built-in IGV browser (Figure 1H). This example showed the exciting power of big data in providing novel insights and quickly developing robust, testable hypotheses with no experimental cost.

In summary, PPRD is a convenient, web-accessible, user-friendly RNA-seq database that allows users to quickly scan the gene expression from maize, rice, soybean, wheat, or cotton public RNA-seq libraries, and returns the multiple forms of results in tables and diagrams, showing the expression levels in various tissues, developmental stages, abiotic stresses, biotic stresses, as well as the differential expression in different mutants and treatments. Our previous Arabidopsis RNA-seq database (ARS) has been updated recently, and the number of libraries has been increased from 20,068 to 28,164 (Zhang et al., 2020). We also plan to continue updating PPRD regularly by including new libraries and new plant species in the future. We believe PPRD will help make the transcriptome bigdata more available and accessible for our plant community members.

## Acknowledgments

We thank all the research groups that contributed RNA-seq data to the public, and we apologize for not being able to cite all the related papers in the main text due to limited space. The list of all public RNA-seq libraries included in the database can be found on the “All Libraries” page. The Program for Guangdong Introducing Innovative and Entrepreneurial Teams (2016ZT06S172).

## Author Contributions

H.Z., Y.Y., Y.L., and Y.S. analyzed the data, H.Z. and Y.Y. processed the data and built the database and website, J.Z oversaw the study. Y.Y., H.Z., and J.Z. wrote the manuscript.

