## Supplementary material for "PPRD: a comprehensive online database for expression analysis of ~45,000 plant public RNA-Seq libraries": Method

**Supplemental Methods**

**RNA-seq data sequencing**

We collected plant public RNA-seq libraries from Gene Expression Omnibus (GEO), Sequence Read Archive (SRA), European Nucleotide Archive (ENA), and DNA Data Bank of Japan (DDBJ) databases using keywords (For example, ‘((Wheat[Organism]) AND "transcriptomic"[Source]) AND "rna seq"[Strategy]’ for wheat). We performed raw reads alignments to respective genomes using HISAT2 (version 2.1.0) (Kim et al., 2015) with parameters (“-max-intron-length= 20000 -k 1 -dta --n-ceil -L,0,0.15” for wheat), and removed duplicated reads using SAMtools rmdup (version 1.4.1) (Li et al., 2009). FPKMs were calculated using Stringtie (version 1.3.3b) (Pertea et al., 2015) with parameters (“-e -r -G”). PPRD also integrates a built-in IGV-Web (https://igvteam.github.io/igv-webapp/) interface to browse and compare the genome-matched RNA-Seq in one library or project. We provided the download button for users to easily acquire the FPKMs of all libraries and detailed library information. More details can be found in the “Tutorials” page.

**Differential gene expression analysis**

Library category was determined into tissue specificity, developmental stages, and stress-related conditions as previously described (Zhang et al., 2020). We manually annotated mutant-related and stress-related libraries as well as their corresponding wild-type and control libraries. Then, differential expression analysis was performed for groups of mutants and treatments using DESeq2 (version 1.32.0) (Love et al., 2014). The top10 groups with the largest differentially expressed genes for mutants and treatments were displayed on PPRD. In addition, the gene expression levels in the top10 groups of different tissues, developmental stages, abiotic stresses, and biotic stresses were also shown on our website.

**Coexpression analysis**

The R package WGCNA (version 1.69) (Langfelder and Horvath, 2008) was utilized to perform weighted gene co-expression analysis. We filtered the libraries with low-quality and low coverage to reduce data noise as previously (Zhang et al., 2020). Then remaining libraries were used to construct co-expression network. In brief, we obtained the optimal soft threshold power for each species based on the scale-free topology criterion. Then, the function blockwiseModules in WGCNA package was used to construct the weighted gene co-expression network and identify the modules with following parameters “TOMType = "unsigned", networkType = "unsigned", maxBlockSize = 40,000, minModuleSize = 30” (Zhang et al., 2020).

Kim, D., Langmead, B. and Salzberg, S.L. (2015) HISAT: a fast spliced aligner with low memory requirements. *Nat Methods* **12**, 357-360.

Langfelder, P. and Horvath, S. (2008) WGCNA: an R package for weighted correlation network analysis. *BMC bioinformatics* **9**, 559.

Li, H., Handsaker, B., Wysoker, A., Fennell, T., Ruan, J., Homer, N., et al. (2009) The Sequence Alignment/Map format and SAMtools. *Bioinformatics (Oxford, England)* **25**, 2078-2079.

Love, M.I., Huber, W. and Anders, S. (2014) Moderated estimation of fold change and dispersion for RNA-seq data with DESeq2. *Genome Biol.* **15**, 550.

Pertea, M., Pertea, G.M., Antonescu, C.M., Chang, T.C., Mendell, J.T. and Salzberg, S.L. (2015) StringTie enables improved reconstruction of a transcriptome from RNA-seq reads. *Nat Biotechnol* **33**, 290-295.

Zhang, H., Zhang, F., Yu, Y., Feng, L., Jia, J., Liu, B., et al. (2020) A comprehensive online database for exploring ∼20,000 public Arabidopsis RNA-Seq libraries. *Mol Plant*.
